# To infect or not to infect: molecular determinants of bacterial outer membrane vesicle internalization by host membranes

**DOI:** 10.1101/763334

**Authors:** Damien Jefferies, Syma Khalid

## Abstract

Outer membrane vesicles (OMVs) are spherical liposomes that are secreted by almost all forms of Gram-negative bacteria. The nanospheres contribute to bacterial pathogenesis by trafficking molecular cargo from bacterial membranes to target cells at the host-pathogen interface. We have simulated the interaction of OMVs with host cell membranes to understand why OMV uptake depends on the length of constituent lipopolysaccharide macromolecules. Using coarse-grained molecular dynamics simulations, we show that lipopolysaccharide lipid length affects OMV shape at the host-pathogen interface: OMVs with long (smooth-type) lipopolysaccharide lipids retain their spherical shape when they interact with host cell membranes, whereas OMVs with shorter (rough-type) lipopolysaccharide lipids distort and spread over the host membrane surface. In addition, we show that OMVs preferentially coordinate domain-favoring ganglioside lipids within host membranes to enhance curvature and affect the local lipid composition. We predict that these differences in shape preservation affect OMV internalization on long timescales: spherical nanoparticles tend to be completely enveloped by host membranes, whereas low sphericity nanoparticles tend to remain on the surface of cells.

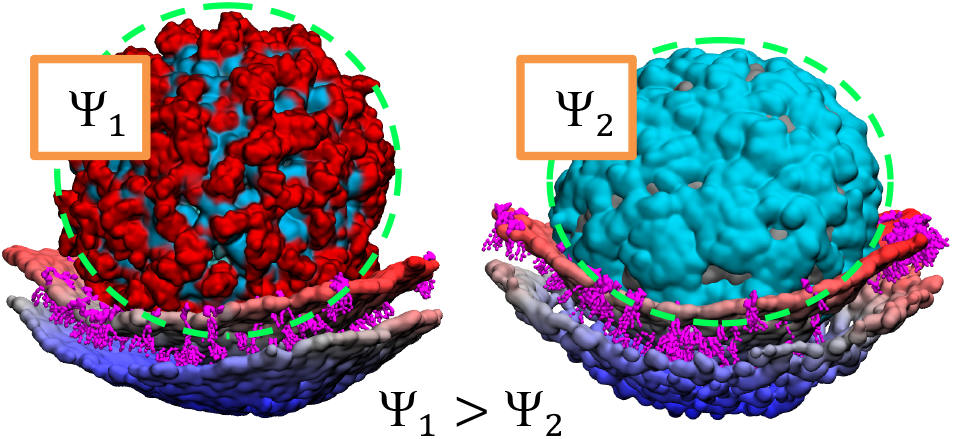

## Introduction

Outer membrane vesicles (OMVs) are membrane-enclosed capsules that are produced by both pathogenic and non-pathogenic Gram-negative bacteria. The nanospheres erupt form parent bacteria either as a stress response, or more commonly, as platforms that are intentionally engineered to mediate essential roles *in vivo.*^1–2^ OMVs have an asymmetric architecture: the extracellular leaflet has a significant proportion of lipopolysaccharide (LPS) macromolecules and the intracellular leaflet has a significant proportion of glycerophospholipids; additional proteins are added to prime OMVs for nutrient scavenging, intercellular communication, producing bacterial biofilms, and mediating bacterial pathogenesis.^3–6^

The abundance of adhesins, toxins, and immunomodulatory compounds are regulated by parent cells to functionalize OMVs for delivering pathogenic cargo and transmitting infectious disease. Adhesins promote initial interactions with epithelial and myeloid cells, immunomodulatory compounds elicit potent immune responses, and toxins break down host cell defenses and attack intracellular organelles. Disease proliferation is stepwise: the nano-sized (20–300 nm) proteoliposomes bud from parent bacteria, traverse the external milieu, latch on to host cell surfaces, and subsequently pass through the peripheral plasma membrane of target cells to invade the cellular cytosol and ravage the intracellular space.^7–10^

Vesicular uptake at the host-pathogen interface is inexorably linked to the length of LPS macromolecules in the outer leaflet.^11^ OMVs can exploit rapid, lipid-mediated uptake pathways when they contain LPS lipids with terminal O-antigen chains (smooth-type); OMV uptake is slower and less efficient when OMVs contain shorter (rough-type) LPS macromolecules. We have already shown that O-antigen chain interactions can collectively increase membrane mechanical strength,^12^ and it has been shown elsewhere that endocytosis is facile for rigid nanospheres and stiff liposomes.^13–16^ Taken together, we infer that smooth OMVs and more adept (than rough OMVs) at wrapping host membranes for endocytosis due to their enhanced mechanical strength; we seek empirical evidence to validate this assumption.

We conducted a coarse-grained molecular dynamics simulation study with the Martini force field^17–19^ to rationalize OMV uptake at the host-pathogen interface. Simulations were performed to analyze the properties of both smooth, and rough OMVs at the host-pathogen interface. The simulations reveal that OMV morphology depends on LPS lipid length: smooth OMVs maintained high sphericity when they interacted with host plasma membrane models, whereas rough OMVs lost their spherical shape when they contacted host cell surfaces. This insight alone helps to rationalize differences in the uptake of smooth and rough OMVs at the host-pathogen interface. Rigid nanospheres are adept at slowly wrapping host membranes for complete encapsulation, whereas lower sphericity nanoparticles tend to rapidly generate large curvatures at the spreading front that makes late stage endocytosis processes more energetically demanding.^20–21^ The simulations additionally reveal that membrane composition impacts host membrane wrapping interactions. Ganglioside head groups acted as a zipper to mediate strong OMV-host cell adhesion and force the plasma membrane around the arched edge of the OMVs. At the same time, the sequestered ganglioside lipids tended to form aggregates within the host membranes that (i) promoted local bilayer bending, and (ii) increased the local abundances of lipids associated with membrane rafts and endocytosis.^22–27^

## Results and Discussion

Two types of OMVs were studied, smooth and rough (Figure 1A-B). The LPS molecules in the smooth OMVs incorporated 4 units of the *E. coli* O42 O-antigen chain, whereas in the rough OMVs, the LPS molecules did not contain any O-antigen units, full details are given in the supplementary material. For both OMV types, the outer leaflets were composed of LPS and 1-palmitoyl-2-oleoyl-phosphatidylethanolamine (POPE) lipids in a 1:1 ratio; the inner leaflets were composed of POPE and 1-palmitoyl-2-oleoyl-phosphatidylglycerol (POPG) lipids in a 9:1 ratio. Both types of OMVs had diameters (2*r*_*m*_) of 20 nm (based on the position of the hydrophobic core midplane). We note here that this is the same size as the smallest OMVs known to be produced by Gram-negative bacteria, therefore our simulation system size was comparable to the *in vivo* scenario.^3^

**Figure 1.**
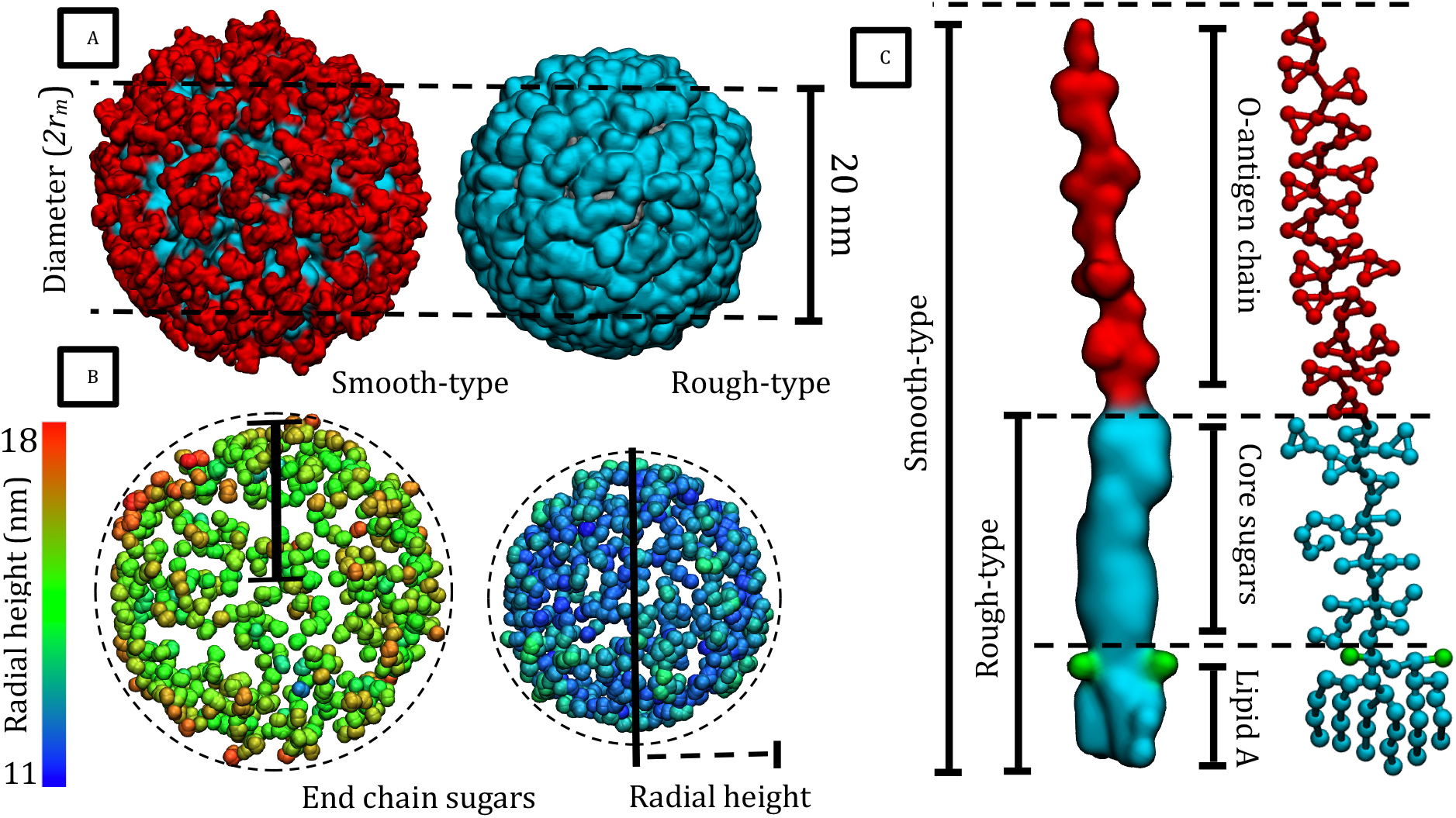
(A) Smooth and rough OMVs—atoms are represented using a volumetric density map. POPE and POPG lipids are silver, LPS molecules have the color scheme: lipid A and core sugars (cyan), O-antigen chain (red). (B) Terminal sugar particles are assigned a BGR color based on their radial height (extension). (C) LPS molecule partitioned into constituent lipid A anchor, core sugar domain, and terminal O-antigen chain. The lipid A phosphate groups are colored green to clarify the position of the water-lipid interface that is referenced throughout. Atoms are represented using a volumetric density map (left), and a simpler ball-and-stick model (right).

### OMVs in water

It is instructive to first provide the lipid packing parameters for the OMVs when they were simulated in water. Parameters were computed using 1 μs long simulations. We stress here that the water-lipid interface was characterized by measuring the radial height of the lipid A phosphate groups (Figure 1C), whereas the conformations of the saccharides were characterized by measuring the full extension of the LPS lipids, i.e. the radial length of the molecules with respect to the center of the OMVs (Table 1). Mean areas per lipid were calculated to quantify lateral packing parameters for each type of molecule in the OMVs. The LPS molecules in the OMV were more tightly packed (less area per lipid), than those from comparable simulations of flat bacterial cell membranes (Figure S1).^28–29^ Despite their longer length, the LPS molecules in the OMVs achieved lateral packing comparable to the shortest LPS (Re-type) lipids found in flat *E. coli* outer membranes.^28–30^ This unusually tight packing of LPS lipids can be ascribed to (i) additional fringe volume that affords LPS head groups more conformational freedom and the capacity to achieve unusually tight lamellar alignment, and (ii) compensatory lateral area expansion of neighboring phospholipids, which pushes the LPS molecules closer together than in flat membranes.

**Table 1:** Summary of lipid properties for OMVs in solution; data are shown for the inner leaflet (IL) and the outer leaflet (OL). Standard deviations are less than 0.02 for areas per lipids and less than 0.9 for radial heights.

| Membrane type | Mean area per lipid (nm<sup>2</sup>) |  |  |  | LPS radial height (nm) |  |
| --- | --- | --- | --- | --- | --- | --- |
|  | LPS | POPE (IL) | POPE (OL) | POPG | phosphates | extension |
| Smooth OMV | 1.57 | 0.55 | 1.26 | 0.60 | 11.63 | 15.38 |
| Rough OMV | 1.46 | 0.50 | 1.20 | 0.53 | 11.52 | 12.42 |
| Flat outer membrane | Re-LPS: 1.59 | 0.62 | — | 0.61 | — | — |
|  | Rough LPS: 1.78 |  |  |  |  |  |
|  | Smooth LPS: 1.90 |  |  |  |  |  |

There were large grooves between the O-antigen chain clusters on the surface of the smooth OMV, whereas the rough OMV surface was relatively smooth and uniform. The differences in surface topology can be explained using simple geometric considerations (see the supplementary material and Figure S2) if we acknowledge (i) the different radial lengths (extension) of the LPS lipids: 12.42 nm (rough LPS), and 15.38 nm (smooth LPS), and (ii) that spherical surface area scales as the square of radial length. The surface of the smooth OMV is reminiscent of the simian vacuolating virus 40 outer edge, which is littered with grooves that bind ganglioside lipid receptors in host cell plasma membranes to mediate host membrane wrapping interactions.^31–34^

There were cohesive intermolecular interactions between terminal O-antigen chains across the entire surface of the smooth OMV, while the rough OMV was held together with fewer cohesive carbohydrate-carbohydrate interactions. The ability of smooth LPS molecules to form clusters has previously been shown in simulations of flat membranes.^28–30^ Since we have already shown that O-antigen chain interactions increase the mechanical strength of (flat) Gram-negative membranes in previous works, it is reasonable to extrapolate that smooth OMVs are stiffer than similarly sized OMVs composed of rough LPS.^12^

### OMVs interacting with simple phospholipid bilayers

In order to obtain a baseline characterization of the interaction of OMVs with simple membranes, the OMVs were simulated with phospholipid bilayers containing only palmitoyl-2-oleoyl-glycero-3-phosphocholine (POPC) lipids (Figure 2A-B). The following simulation data are summarized in Table 2: the axis components of the radius of gyration, the radial heights of the LPS lipids, the number of POPC lipids within 5 nm of the OMVs, and the percentage of LPS molecules bound to POPC lipids after 2 microseconds. Comparable to elastic lipid-covered nanoparticles,^13^ the rough OMV deformed and lost its spherical shape when it interacted with the host lipid membrane. The OMV distorted and spread out along the membrane surface—an event that tends to precede the incomplete wrapping of host membranes in comparable computer simulations.^13–16^ In addition to this large-scale shape loss, there was deformation of individual LPS lipids: compared with the simulation in water, there was compression of the rough LPS lipids along their long axis.

**Figure 2.**
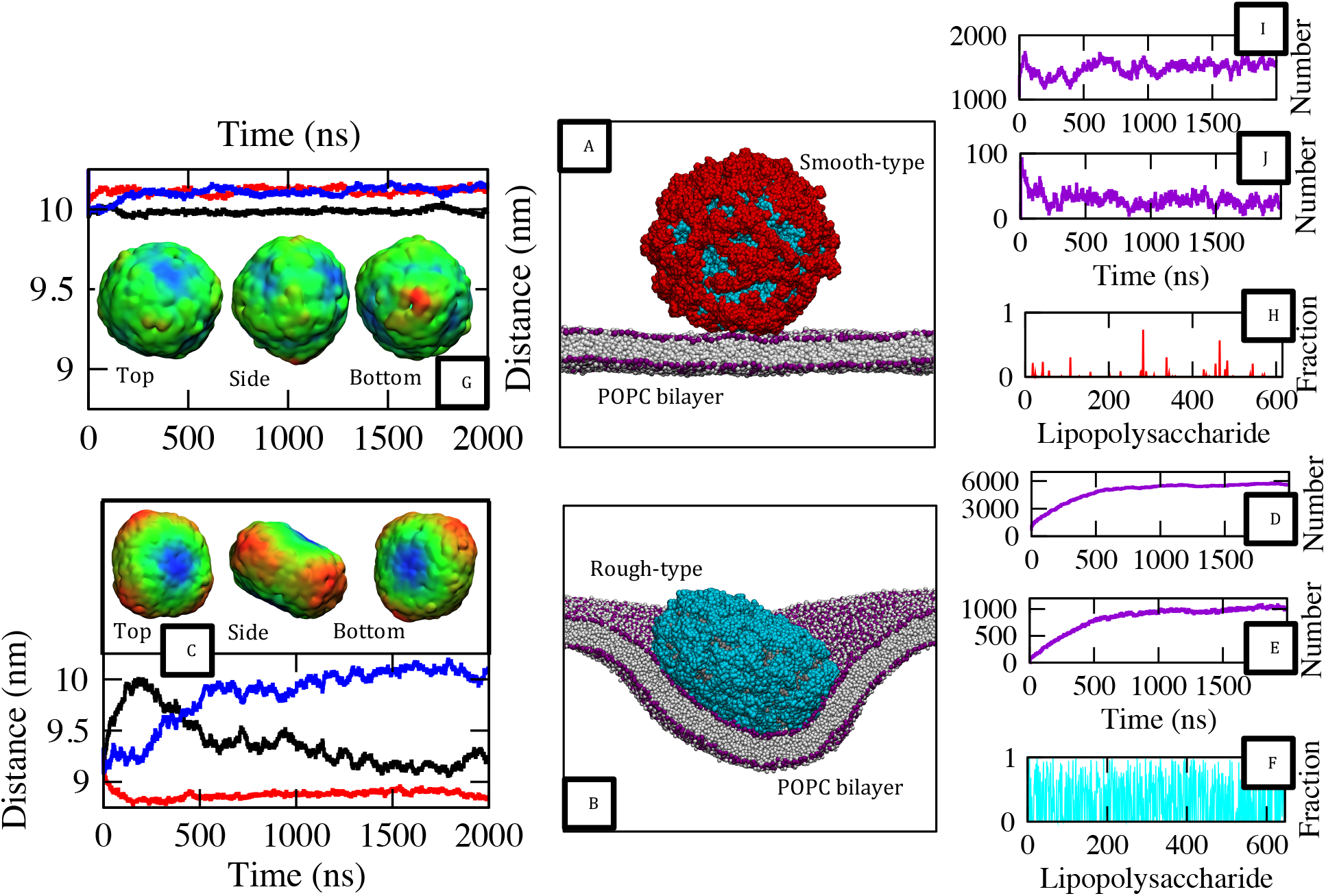
(A-B) Smooth and rough OMVs at the POPC bilayer. (C) Axis components of the radius of gyration for the rough OMV (bottom); phosphate group (BGR) height map after 2 μs (top). (D) POPC lipid shell population for the rough OMV (5 nm cutoff); (E) LPS-POPC contact number (0.6 nm cutoff). (F) Fraction of simulation frames—during the last 0.25 μs—with registered LPS-POPC contacts (per rough LPS molecule). (G) Axis components of the radius of gyration for the smooth OMV (top); phosphate group (BGR) height map after 2 μs (bottom). (H) Fraction of simulation frames—during the last 0.25 μs—with registered LPS-POPC contacts (per smooth LPS molecule). (I) POPC lipid shell population for the smooth OMV; (J) LPS-POPC contact number. Figures I-J, and Figures D-E are measured from the point of OMV-host membrane first contact and thus, are non-zero from the start.

**Table 2:**
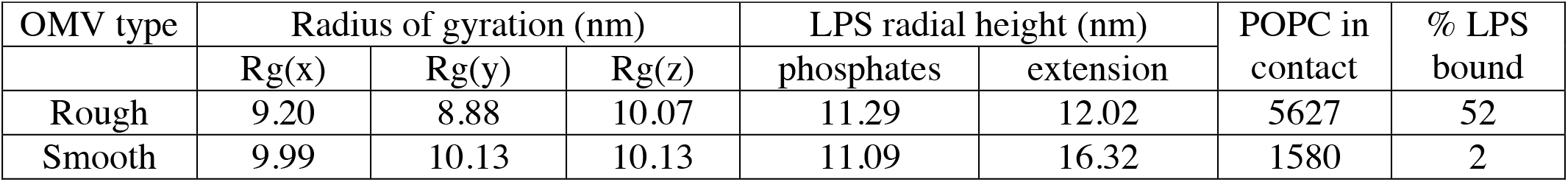
Summary of OMV properties at the POPC bilayer. Standard deviations for radius of gyration values are less than 0.07 and for heights they are less than 1.74

The POPC bilayer also deformed as it wrapped around the oblate OMV. We state here that the number of POPC lipids in contact with the OMV was used as a measure of membrane wrapping around the OMV. After 2 microseconds, there were over 5000 POPC lipids within 5 nm of the rough LPS lipids (Figure 2D), and over 50% of the rough LPS molecules were bound to POPC molecules (Figure 2E-F). The correlation coefficient for membrane wrapping with respect to sampled simulation time was negative (−0.55) during the last 250 nanoseconds, indicating that membrane wrapping peaked during simulation time. Further OMV internalization seems unfeasible based on (i) this late stage reduction in host membrane wrapping, (ii) the loss of rough OMV sphericity, and (iii) comparisons with large-scale simulations, and theories of membrane wrapping around nanoparticles.^13–16^

In stark contrast, the smooth OMV maintained high sphericity throughout simulation time, and did not induce any appreciable wrapping of the POPC membrane upon interaction. We note here that stiff elastic nanoparticles that wrap host membranes are expected, according to theoretical analysis, to promote gradual areal deformation and bilayer bending.^14–16^ Voronoi tessellation analysis of the phospholipid membrane revealed that there was a disparity in lipid packing between the POPC bilayer leaflets at the end of the simulation: the upper leaflet had an area per lipid value of 0.54 nm^2^— ordinarily 0.68 nm^2 21^—and the lower leaflet had an area per lipid value of 0.81 nm^2^. The lipid packing was relatively uniform per bilayer leaflet (Figure S3), indicating that the whole bilayer was strained, rather than there being local regions of high strain close to the OMV. Despite the disparity in upper and lower leaflet lipid packing, and the evident bilayer strain, there was a lack of host membrane wrapping around the OMV: no more than 2% of the smooth LPS molecules were in contact with POPC lipids during the last 250 nanoseconds (Figure 2H) and there were only 1580 POPC molecules within 5 nm of the LPS molecules for the endpoint conformation (Figure 2I-J). Contacts between LPS and POPC molecules were short-lived, the POPC lipids were equally mobile in the upper and lower leaflets: lateral diffusion coefficients were 0.063 ± 0.0003 1×10^−5^ cm^2^/s in both leaflets. Taken together, it is apparent from these data that the bilayer was strained and there were some forces favoring membrane wrapping, but the impermanent LPS-POPC interactions, coupled with the unfavorable energy associated with perturbing the stable bilayer structure, hindered the POPC lipids engulfing the OMV to any appreciable extent on the timescale of these simulations.

### OMVs interacting with model plasma membranes

Given that ganglioside head groups support strong cohesive interactions with invasive toxins, and tend to facilitate membrane reshaping due to their high intrinsic positive curvature, we next explored the interaction of OMVs with more complex models of host membranes.^36–41^ Both OMVs were simulated with a pre-equilibrated,^32–45^ multicomponent plasma membrane model composed of POPC (25%), POPE (25%) 1-palmitoyl-2-oleoyl phosphatidyl serine, or POPS (7.5%), GM3 ganglioside (5%), sphingomyelin (7.5%), cholesterol (25%) and phosphatidylinositol 4,5-biphosphate, or PIP_2_ (5%). Table 3 and Table 4 summarize the findings from these simulations, which are described in detail below.

**Table 3:** summary of OMV properties at the host plasma membrane. Standard deviations for radius of gyration values are less than 0.01 and for heights they are less than 1.7.

| OMV type | Radius of gyration (nm) |  |  | LPS radial height (nm) |  |
| --- | --- | --- | --- | --- | --- |
|  | Rg(x) | Rg(y) | Rg(z) | phosphates | extension |
| Rough | 9.83 | 8.76 | 9.43 | 11.22 | 11.99 |
| Smooth | 10.35 | 10.17 | 10.12 | 11.21 | 14.18 |

**Table 4:** Changes in local lipid composition within 5 nm of the OMVs. Data are shown for each lipid type (per bilayer leaflet).

| OMV type | Enrichment/depletion within 5 nm of the OMV (%) |  |  |  |  |  |  |  |  |  |  |  |  |  |
| --- | --- | --- | --- | --- | --- | --- | --- | --- | --- | --- | --- | --- | --- | --- |
|  | GM3 |  | PIP<sub>2</sub> |  | Cholesterol |  | Sphingolipid |  | POPE |  | POPC |  | POPS |  |
|  | IL | OL | IL | OL | IL | OL | IL | OL | IL | OL | IL | OL | IL | OL |
| Rough | — | 2.0 | 2.3 | — | 12.6 | 1.5 | — | 7.3 | 2.0 | -11.2 | -12.1 | -1.1 | 6.0 | — |
| Smooth | — | -2.0<sup>†</sup> | 18.4 | — | 18.6 | 7.9 | — | 4.5 | -14.7 | -10.2 | -12.0 | -4.0 | 3.9 | — |

<sup>†</sup> There was an absolute increase in the number of GM3 gangliosides (within 5 nm), but a relative reduction in GM3 concentration, given the non-negligible influx of sphingomyelin and cholesterol molecules into the local OMV ‘shell’.

The rough OMV lost its spherical shape when it interacted with the model plasma membrane: the axis components of the radius of gyration were more than 10% different for the vesicle after 2 μs. The LPS lipids were found to compress towards the center of the OMV as it spread over the membrane surface. For the last 50 nanoseconds there were 6575 lipids within 5 nm of the OMV and 14% of the LPS molecules were in contact with ganglioside lipid head groups (based on a 0.6 nm cutoff), showing that the membrane had partially wrapped around the OMV to form a ‘pit’ or invagination. The height of the membrane pit was computed by exporting the *z*-axis coordinates of host membrane phosphate groups within 5 nm of the OMV. Based on this procedure, the height range (*z*-axis) for the plasma membrane pit was 19.1 nm after it had interacted with the multicomponent plasma membrane, compared with 21.0 nm for the POPC bilayer simulation. In other words, the deformation of the rough OMV and the host membranes which they bind, were similar regardless of specific host membrane composition. Comparable results were obtained when soft nanoparticles were simulated with flexible membranes: deformation was partitioned between the soft nanospheres and the flat bilayers due to their low flexural moduli.^14–15^

The smooth OMV retained its spherical shape after it interacted with the model plasma membrane, comparable to the POPC bilayer simulation we discussed previously (Figure 3A, 2A). There was slight asymmetric perturbation of the Lipid A phosphate plane, which defines the position of the water-lipid interface in this paper (Figure 3B). The phosphate groups had a radial height of 11.21 ± 0.57 when the OMV was at the surface of the model plasma membrane, compared with 11.63 nm in water. The average length of the smooth LPS was reduced to 14.18 ± 1.00 nm after the OMV interacted with the host plasma membrane (Figure 3C), compared to 15.4 nm in water and 16.3 nm when interacting with the POPC bilayer. This reduction in the radial extension of smooth LPS is indicative of rather different interactions with the lipids of the plasma membrane compared to molecular interactions with both the simple membrane and just water, and therefore it is instructive to consider these interactions in detail.

**Figure 3.**
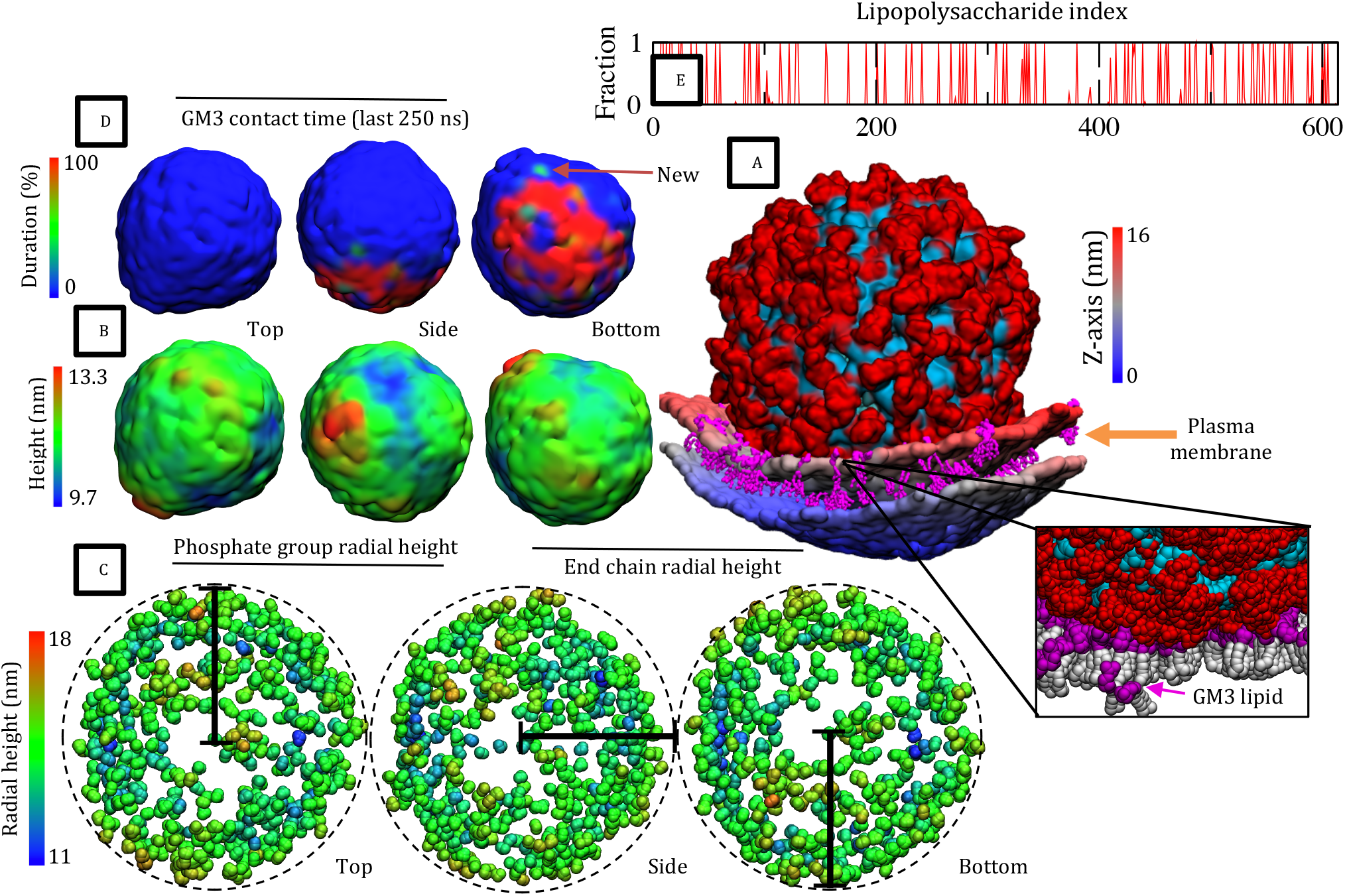
(A) Smooth OMV and the host plasma membrane (5 nm shell). Phosphate groups are assigned a (BSR) color based on their height (*z*-axis coordinate); ganglioside molecules are magenta. In the inset, ganglioside head groups are magenta; lipid tails are white. (B) Phosphate group (BGR) height map after 2 μs. (C) Terminal sugar particles are assigned are a (BGR) color based on their endpoint radial height. (D) LPS-GM3 contact duration projected onto the phosphate group density map; data are shown for the last 0.25 μs of simulation time. (E) Fraction of simulation frames— during the last 0.25 μs—that LPS-GM3 contacts were registered for each one of the 615 smooth LPS molecules in the OMV outer leaflet. Put simply, the graph shows the fraction of sampled simulation time that each smooth LPS lipid was bonded to GM3 lipids (based on a 0.6 cutoff). Once LPS-GM3 interactions were formed, they were almost always maintained thereafter.

The flexible O-antigen chains of the smooth LPS molecules changed their packing as the OMV interacted with the carbohydrate head group moieties of the GM3 lipids. After 2 microseconds, 15.6 % of the OMV LPS molecules were in contact with GM3 lipids, and the interactions between O-antigen chains and ganglioside lipids were long-lived: once formed, the interactions were almost always maintained thereafter (Figure 3D-E). GM3 lipids traversed the length of the plasma membrane and sporadically migrated towards the smooth OMV. The domain-favoring ganglioside lipids^41,46–47^ became trapped within the vicinity of the OMV (Figure S4) and increasingly contributed to local bilayer perturbation. The GM3 lipids affected local lipid packing, lipid diffusion and membrane curvature. The lipid arrangement, and the abundance of cohesive GM3– LPS interactions, promoted slow, axisymmetric wrapping of the plasma membrane around the OMV. For the endpoint system configuration at 2 microseconds, the composition of the local bilayer (based on a 5 nm cutoff) had changed e.g. there was a relative increase of PIP_2_ (18.4 %) and cholesterol (18.6 %) in the (inner) intracellular leaflet, and a relative increase of sphingomyelin (4.5 %) and cholesterol in the extracellular leaflet (7.9 %). The changes in bilayer composition can be explained by lipid shape: for example, PIP_2_ has an inverted conical shape that favors the high positive curvature of the expanding inner bilayer leaflet and cholesterol molecules, which are small and taper from membrane core to membrane surface, tend to fill the spaces between these inverse funnel-shaped (PIP_2_) lipids.

The OMV-plasma membrane interaction mimics the docking of simian vacuolating virus 40 (SV40) onto host cell plasma membranes: the LPS lipids interlink domain-favoring ganglioside lipids that promote bilayer curvature and membrane wrapping.^31–34^ The number of GM3 lipids within 5 nm of the OMV was described quite accurately by the generalized logistic function (correlation coefficient of 0.93), which suggests that membrane wrapping could be imperceptibly slow going forward. But based on similarities between our data, and trajectories from large-scale simulations,^13–16^ we reason that OMVs can be primed for complete internalization by increasing the concentration, and length, of smooth LPS lipids (in the outer leaflet) and likely also the length of the ganglioside molecule, i.e. GM1. ^31–34^

### Phospholipid vesicles and model host membranes

To better understand how the LPS outer shell impacts OMV uptake, we investigated how a vesicle behaved at host cell membranes when it was made with POPE and POPG lipids alone. The vesicle contained POPE and POPG lipids in a 9:1 ratio (per bilayer leaflet); it was initially simulated in water (Figure 4A) and was subsequently simulated with the POPC bilayer and the plasma membrane model. The vesicle had a diameter (*2*r*_*m*_*) of 20 nm to make it physically comparable to the smooth and rough OMVs presented earlier. We emphasize here that these simulations were control experiments, they help us to understand how the LPS lipid leaflet affects wrapping interactions through comparative analysis. The simulations were not performed to directly emulate *in vivo* interactions, instead they were conducted to make this research more robust. For clarity, we will first present the lipid packing parameters for the vesicle when it was simulated in water, and we will then present the simulation results for the vesicle when it was simulated with the POPC bilayer in the first instance, and the multicomponent plasma membrane model in the second.

**Figure 4.**
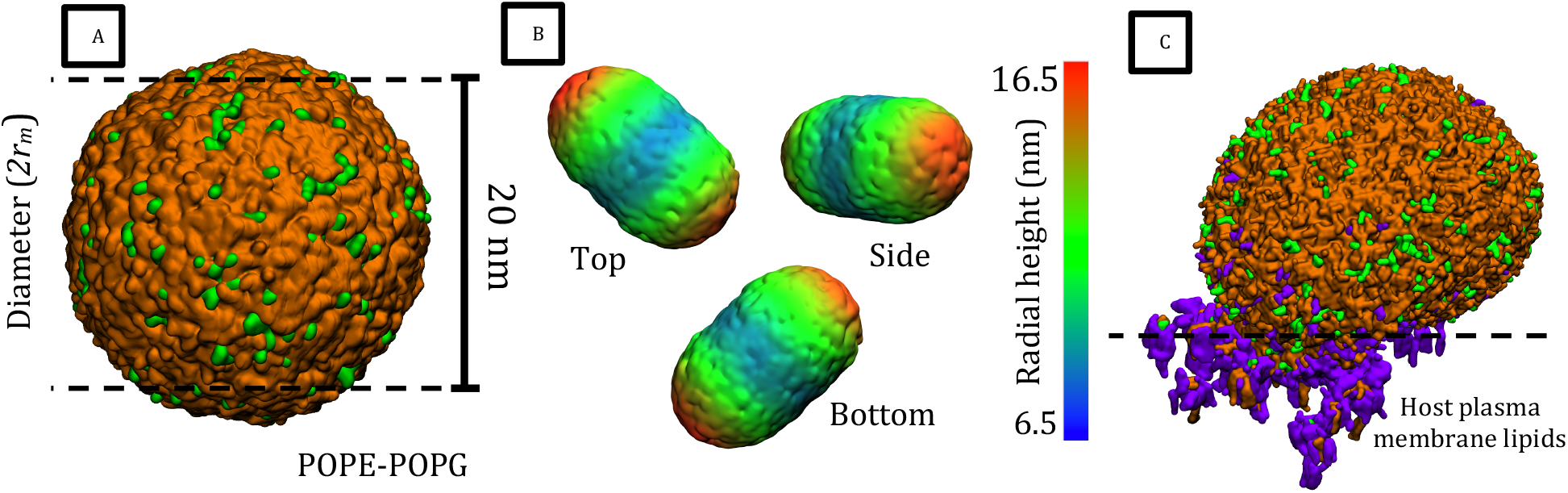
(A) POPE-POPG vesicle used in the control experiments—atoms are represented using a volumetric density map; POPE lipids are orange and POPG lipids are green. (B) Phosphate group (BGR) height map for the POPE-POPG vesicle after it bounces off the host POPC membrane. (C) Endpoint conformation for the simulation of the POPE-POPG vesicle and the multicomponent plasma membrane model. POPE molecules are orange, POPG molecules are green, and the host plasma membrane lipids that are within 0.5 nm of these lipids are purple. The vesicle fuses with the host plasma membrane to form a lipid-lined pore, which facilitates lipid exchange between the plasma membrane and the POPE-POPG vesicle.

When the vesicle was simulated in water, the mean area per lipid values were 0.67 ± 0.001 nm^2^ (POPE), and 0.70 ± 0.01 nm^2^ (POPG), while the membrane thickness was 3.86 ± 0.01 nm. Compared with flat bacterial membranes, the average areas per lipid were increased (12.2 % and 11.4%), and the membrane was 3.3 % thinner.^30.48^ If we compute the areas per lipid for each membrane leaflet it becomes clear that the unusual packing parameters were due to non-uniform (total) surface area along the lipid long-axis. Since the area per lipid increases along the membrane normal, lipid head groups are pushed closer together in the inner leaflet and further apart in the outer leaflet (Table 5). Comparable lamellar organization has already been described in previous simulations that were performed with the Martini coarse-grained force field.^49^

**Table 5:** Areas per lipid for the POPE-POPG vesicle in water. Standard deviations are less than 0.01 throughout.

| Membrane type | Mean area per lipid (nm<sup>2</sup>) |  |  |  |
| --- | --- | --- | --- | --- |
|  | POPE (IL) | POPG (IL) | POPE (OL) | POPG (OL) |
| Vesicle | 0.51 | 0.53 | 0.78 | 0.82 |
| Flat membrane | 0.62 | 0.61 | 0.62 | 0.61 |

The phospholipid vesicle deformed when it was simulated with the POPC bilayer. For the last 0.25 μs, the axis components of the radius of gyration (*x*, *y*, and *z*) had a difference of up to 13.5 %, and the radial heights of the outer leaflet phosphate groups were up to 73 % different (Figure 4B). The vesicle underwent approximate axisymmetric deformation to morph into an eccentric spheroid structure after the vesicle made transient contact (approximately 1 ns) with the POPC bilayer, and subsequently moved away from the membrane surface. This observation reveals that even transient contacts with host membranes can deform the morphology of softer vesicles.

When the vesicle was simulated with the multicomponent plasma membrane model, the lipid head groups of the vesicle merged with the head groups of the plasma membrane at the point of contact. The fusion event was in line with the stalk-pore intermediate model, which explains membrane fusion via a stepwise process that involves hemifusion stalk intermediates.^50^ Comparable to previous works,^51–54^ there was initial membrane aggregation, subsequent membrane association, lipid rearrangement, and membrane content mixing. Apposed head groups merged to form a lipid-lined pore that facilitated the exchange of content from the vesicle to the host plasma membrane, and likewise from the host plasma membrane to the vesicle (Figure 4C).

The widths of hydrophilic head group domains were computed for the OMVs and the POPE-POPG vesicle to better understand why direct membrane fusion was suppressed in certain simulations, and facilitated in others. The width of the peripheral head group domains was determined by computing radial distribution functions for polar and charged Martini type beads with respect to vesicle centers. Based on this metric, the POPE-POPG vesicle head group domain was 0.50 ± 0.25 nm, and the width of the rough and smooth OMV head group domains were 1.78 ± 0.24 nm and 5.01 ± 0.21 nm, respectively. Given that lipid protrusions instigate membrane fusion via pre-stalk transition states, the data suggests that OMVs will be less likely to achieve direct membrane fusion than phospholipid liposomes. Lipid tails in OMVs must circumvent a thicker barrier of hydrophilic head groups to fuse with (hydrophobic) host membrane cores. In other words, OMVs have thick shells of hydrophilic sugars that suppress lipid tail aggregation and membrane core content mixing, whereas glycerophospholipid vesicles are less encumbered by their hydrophilic head group moieties.

We would like to stress here that membrane fusion reactions are generally non-trivial and fusion energy profiles depend on almost all membrane parameters e.g. membrane curvature, lipid heterogeneity, and the abundance of POPE molecules, which have intrinsic negative curvature and consequently, the capacity to stabilize the concave necks of hemifusion intermediates.^55–56^ Indeed, this complexity between membrane parameters and fusion energy profiles is apparent in our simulations: the POPE-POPG vesicle fused with one membrane and not the other. We are not stating that direct membrane fusion reactions are governed by head group width alone, but rather that OMV head group domains are unusually thick; so much so, that OMVs do not fuse with host membranes regardless of simulation setup. Based on these observations, we can conclude that OMVs must enter cells via host membrane wrapping interactions, which favor nanosphere stiffness and strong cohesion forces.

## Conclusion

We demonstrated that the length of LPS lipids affects OMV properties at the host-pathogen interface. When OMVs contain O-antigen chain polymers (smooth-type) they maintain high sphericity at host cell membranes. When OMVs contain shorter LPS lipids (rough-type) they spread out across the host membrane surface—an event that is known to precede the incomplete wrapping of host membranes. The internalization events are also affected by plasma membrane composition: ganglioside lipid receptors act as a zipper to mediate strong OMV-host cell adhesion, which according to theoretical analysis, helps to facilitate complete nanoparticle encapsulation.^14–16^ Additionally, the trapping of ganglioside lipids (by OMVs) tends to increase local bilayer curvature and the local abundances of lipids that are associated with domain formation (cholesterol and sphingomyelin)^22–23^ and endocytosis (PIP_2_).^24–27^ Throughout our simulations, rough OMVs tended to deform host membranes more rapidly than their smooth OMV counterparts: smooth OMVs were associated with slow increases in elastic energy, whereas rough OMVs were associated with faster curvature generation.

The simulations can be used to rationalize differences in the uptake of smooth and rough OMVs at the host-pathogen interface.^11^ The cohesive interactions between terminal O-antigen chains confer mechanical strength to smooth OMVs,^12^ which are robust and tend to retain high sphericity at the host-pathogen interface. When OMVs lack terminal O-antigen chains they are more flexible and tend to spread out across the membrane surface. The differences in OMV rigidity affect host membrane wrapping: rigid nanospheres are adept at slowly wrapping host membranes to minimize the size of the energy barrier that must be overcome during the late stages of nanosphere encapsulation.^14–16^ In contrast, soft nanospheres tend to generate large curvatures at the spreading front that makes nanosphere encapsulation less likely.^20–21^ Smooth OMVs are stiffer than rough OMVs, and consequently, they are more likely to achieve full host membrane wrapping by slowly forcing the bilayer to adhere to their round surface.

It is important to note here that our simulations dispel the common assumption that OMVs readily pass through host membranes by direct membrane fusion.^7–9^ Our simulations demonstrate that membrane fusion is affected by vesicle head group domain width. Phospholipid vesicles had small head groups in our simulations and they readily achieved membrane fusion via the formation of hemifusion stalk intermediates. Conversely, OMVs had thick shells at their outer edge that curbed membrane fusion by impeding membrane aggregation. The distances between host membranes and OMVs were large throughout our simulations, making it difficult for them to fuse. The results suggest that OMVs more commonly enter cells through host membrane wrapping—a process that favors vesicle stiffness. We do not entirely discount direct membrane fusion as an OMV uptake pathway, but we discount the common assumption that direct membrane fusion is facile *in vivo*. Fusion barriers seem to scale with head group width, implying that direct membrane fusion is hardly accessible for OMVs with complete core saccharide section, and even less accessible when they contain terminal O-antigen chains.

Comparable to theoretical studies, we found that membrane wrapping interactions also depend on both cohesion energies and bilayer bending moduli. Smooth OMVs tended to wrap host membranes that contained ganglioside lipids, which lower bending moduli and foster strong cohesive interactions, but were unable to change to shape of POPC membranes, which have higher bending moduli and do not foster cohesive carbohydrate interactions. Strong cohesion forces were generated through tight interlinking of LPS ligands and ganglioside receptors that helped the plasma membrane to adhere to the arched edge of the smooth OMV. The OMVs sequestered gangliosides within the host plasma membrane to produce large ganglioside clusters, which have been found to lower local bending parameters and dramatically enhance preferences for spontaneous curvature and bilayer reshaping.^39–40,57–58^ Plasma membrane heterogeneity also affected bilayer bending: constituent lipids were able to reorganize themselves to reduce line tension energies e.g. PIP_2_ lipids were preferentially partitioned to the inner plasma membrane leaflet when it was made round through interactions with the smooth OMV. Lipid type heterogeneity was well established as a driving force for curvature generation in experimental works,^59–61^ but it is also evident in other coarse-grained molecular dynamics simulation studies that were conducted with the Martini force field.^42^

The uptake of OMVs is multifaceted and quite complex; lipid composition affects the stiffness of both invasive OMVs and flat bilayers, and at the same time, the strength of the cohesive forces that are generated when they interact with each other. Further, there is dynamic lipid co-clustering associated with OMV docking that affects how easy it is for OMVs to wrap host membranes. For clarity, we have expressed interesting simulation results schematically. The figure affords some insights into the membrane modulating effects of smooth OMVs in our simulations and also provides a more general hypothesis for OMV uptake (Figure 5). We found that LPS lipids, much like cholera and Shiga toxin,^37,62^ have the capacity to modulate bilayer properties by binding to, and rearranging, ganglioside lipids within host cell membranes. Comparable to stiff icosahedral SV40 virions,^31–34^ smooth OMVs initially bind ganglioside lipid receptors in host membranes and subsequently create a hemispherical pit, which can be considered a membrane budding intermediate.^63–64^ The process unfolds as ganglioside molecules stochastically traverse the host membrane surface and sporadically come into contact with the anchored OMV. The ganglioside lipids become trapped at the OMV contact edge, and increasingly affect local lipid packing and bilayer bending moduli. Going forward, it is likely that the host membrane would then be wrapped around the smooth OMV on long timescales; the precise degree of wrapping would be influenced by flexural moduli, cohesion energies, and specific lipid compositions.

**Figure 5.**
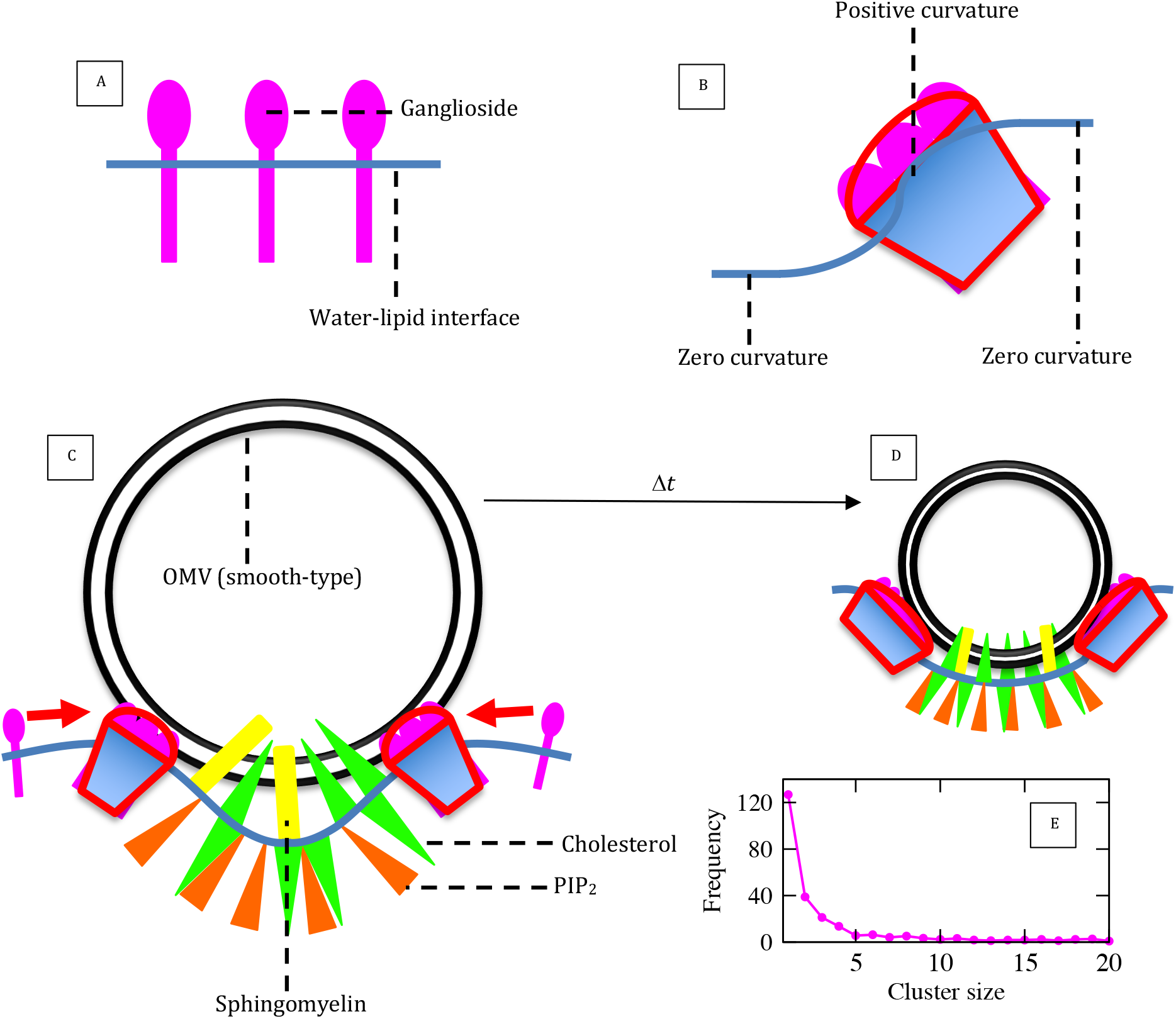
(A) Ganglioside molecules (pink), which are confined to the upper (extracellular) leaflet, create non-negligible stress in plasma membranes that favors bilayer curvature. (B) Energy barriers for bilayer reshaping are reduced when ganglioside molecules aggregate and form clusters that have high intrinsic positive curvature. Ganglioside lipid aggregates can reduce line tension between flat and warped bilayer domains due to their conical shape. (C-D) Schematic illustrations showing how smooth OMVs affect bilayer shape and composition; sphingomyelin are yellow rectangles, PIP_2_ lipids are orange triangles, cholesterol molecules are inverted green triangles; POPS, POPC, and POPE lipids are omitted for clarity. After simulation time (Δ*t*), a significant number of (GM3) gangliosides have interlinked the smooth LPS lipid head groups (based on a 0.6 nm cutoff) and consequently, there is a change in local lipid composition and bilayer curvature. (E) Abundance of ganglioside monomers and aggregates detected (during the last 100 ns of simulation time) when the plasma membrane was simulated without OMVs; ganglioside aggregation is discussed more in the supplementary section.

From this, it is clear that OMVs can be primed to fully wrap host membranes by increasing the concentration, and length, of smooth LPS lipids in the outer leaflet. The modifications will increase OMV stiffness and make lipid-mediated uptake pathways more accessible—explaining why smooth OMVs with longer O-antigen chains can more efficiently exploit lipid-mediated invasion pathways in experimental studies.^11^ We can also infer from our simulations, and theories of nanosphere encapsulation, that the composition of host membranes can be manipulated to suppress OMV encapsulation. Membrane bound biomolecules change the lipid distribution and stiffness constants of the bilayers that they are embedded in. For example, integral membrane proteins can make host membranes stiffer and can also affect ganglioside co-clustering and ganglioside lipid distribution.^45^ Stiffer host membranes that have fewer unconstrained ganglioside head groups would be much less prone to wrap around invasive OMVs and lipid-mediated uptake events would be suppressed. In other words, OMV uptake efficiency *in vivo* would depend on resident host cells, given that membrane composition differs for cells throughout the human body.^65–67^ Our simulations were by no means comprehensive, but they were sufficiently diverse to resolve an interesting question: why does OMV lipid-mediated uptake depend on the cell wall architecture of parent (bacterial) cells?^11^ The coarse-grained simulations answer this question, and in doing so, makes some headway in clarifying longstanding uncertainties associated with OMV uptake e.g. why does cholesterol, Filipin, and caveolae concentration affect OMV internalization? And why does LPS modification affect immunogenicity or the efficacy of OMV vaccine adjuvants?^8^

## Supporting Information

Molecular dynamics simulation details; simulation analysis details; discussion of OMV surface topology; details of supporting simulations that are not directly germane to this communication; discussion of ganglioside lipids and ganglioside clustering; limitations.

## Acknowledgments

For the provision of computational resources, we thank (i) the HECBioSim Consortium, and (ii) The University of Southampton High Performance Computing Facility.

